# Effect of alleviating fibrosis with EGCG-modified bone graft in murine model depended on less accumulation of inflammatory macrophage

**DOI:** 10.1101/2020.11.10.376590

**Authors:** Dengbo Yao, Song Jin

## Abstract

In response to current trends in the modification of guided bone regeneration (GBR) materials, we aimed to build upon our previous studies on epigallocatechin-3-gallate (EGCG) by immersing a commonly used bone graft primarily composed of hydroxyapatite (HA) in EGCG solution, expecting to obtain superior bone–material integration after implantation. Bone grafts are commonly used for bone repair, in which the bone extracellular matrix is stimulated to promote osteogenesis. However, due to its pro-fibrosis effect, this osteoconductive material commonly exhibits implant failure. In addition to providing a basic release profile of EGCG-modified bone graft (E-HA) to clarify the relationship between this material and the environment, we have examined the integration effect via subcutaneous implantation experiments. In this manner, we have assessed the aggregation of pro-inflammatory macrophages, initial angiogenesis, the formation of fibrous capsules, and an enhanced cell viability observed in cultured RAW 264.7 cells. Among these results, we focus on pro-inflammatory macrophages due to their close relationship with fibrosis, which is the most important process in the immune response. Immunofluorescent staining results showed that E-HA substantially compromised the formation of fibrous capsules in hematoxylin-eosin-stained sections, which exhibited less pro-inflammatory macrophage recruitment; meanwhile, the cell viability and primary angiogenesis were improved. This work lays the foundation for future studies on GBR.

## 1 Introduction

In guided bone regeneration (GBR), it is important to prevent the damaged area from being filled with soft tissue, leaving no space for bone tissue deposition. This aim can be attained by removing non-osteogenic tissues that interfere with bone regeneration while implanting biomaterials. The design of implanted bone substitutes should be based on the characteristics of the defect; moreover, these substitutes should restore the tissue microenvironment to its pre-impairment state and avoid severe fibrosis to promote integration and function recovery^1,2^. As a primary component of mineral bone, hydroxyapatite (HA), which constitutes 60%–70% of bone and 98% of dental enamel, exhibits good biocompatibility, high osteoconductivity^3^, and a slow absorption rate at the implant site^4^. Accordingly, HA is one of the most widely used bone materials in clinical practice. In a 20-year follow-up study of maxillary sinus floor elevation based on commercial bovine-derived bone mineral, which is primarily composed of HA, the new mineralized bone volume remained stable and even increased from 16.96% to 22.05% over 20 years, while the volume of the implanted graft decreased from 35.87% to approximately 4%^5^. However, in some cases, HA implantation has resulted in disappointing or unsuccessful healing outcomes with thick fibrous capsules, which can inhibit bone–biomaterial integration via fibrosis. In addition, the formation of fibrous capsules is closely related to the host immune response, which has informed the design of next-generation GBR membranes and filling grafts based on a modulated immune microenvironment. With respect to biocompatibility, degradation rate, and mechanical support, the current mainstream clinical needs are no longer being met^6–8^. Epigallocatechin-3-gallate (EGCG), the main catechin of tea, has been considered as a useful substance for modifying HA, due to its multiple therapeutic effects on various human diseases via its anti-inflammatory^9^, anti-carcinogenic^10^, anti-microbial, and anti-oxidative ability^11^. In addition, in the field of bone remodeling, EGCG can promote osteogenesis with increased expression of bone morphogenetic protein-2 (BMP-2), runt-related transcription factor-2 (Runx-2), alkaline phosphatase (ALP), osteonectin, and osteocalcin. Thus, superior ALP activity and bone defect mineralization of bone defect can be achieved^12^ to prevent inflammatory bone loss caused by inhibited prostaglandin E synthesis^13^. In a previous study, we found that the addition of EGCG to a designed material introduced a promising microenvironment for bone regeneration, based on macrophage recruitment and phenotype^14^. In macrophages, the persistence of inflammatory phenotypes can contribute to fibrosis, which impacts the overall bone growth. Although the mechanism of fibrosis that occurs in implant biodegradation and the blocking interaction between the biomaterial and surrounding tissue require further study, it has been confirmed that fibrosis is associated with the participation of certain macrophage phenotypes and various immune cells. The fibrosis response usually begins with a severe foreign body reaction (FBR) as cells are damaged, necrotized, and stressed^15–17^. Nevertheless, principal cells that promote tissue repair are no different from those leading to fibrous capsules, except for the persistence and drastic communication of inflammatory cells and myo-fibroblasts^18^. In fact, with different degrees of fibrosis, the outcome of implantation can be beneficial for repair and prognosis (with the deposited provisional extracellular matrix [ECM] providing a frame for angiogenesis) or harmful to the host (impairing the integration and resulting in failure)^19^. Hence, the ideal situation is to avoid serious FBR and to enable direct healing by controlling the infiltrated cells and their ability to modulate the immune response. In summary, immunoreactions play a vital role in the healing process and determine the final result. Considering the broad clinical application of HA, it is valuable to understand the immune response invoked by the involvement of EGCG, which can reduce the level of pro-inflammatory macrophages. The purpose of the present study was to evaluate whether binding EGCG to HA can mitigate FBR with less fibrosis, thus enhancing bone regeneration (**Fig. 1**).

**Figure 1.**
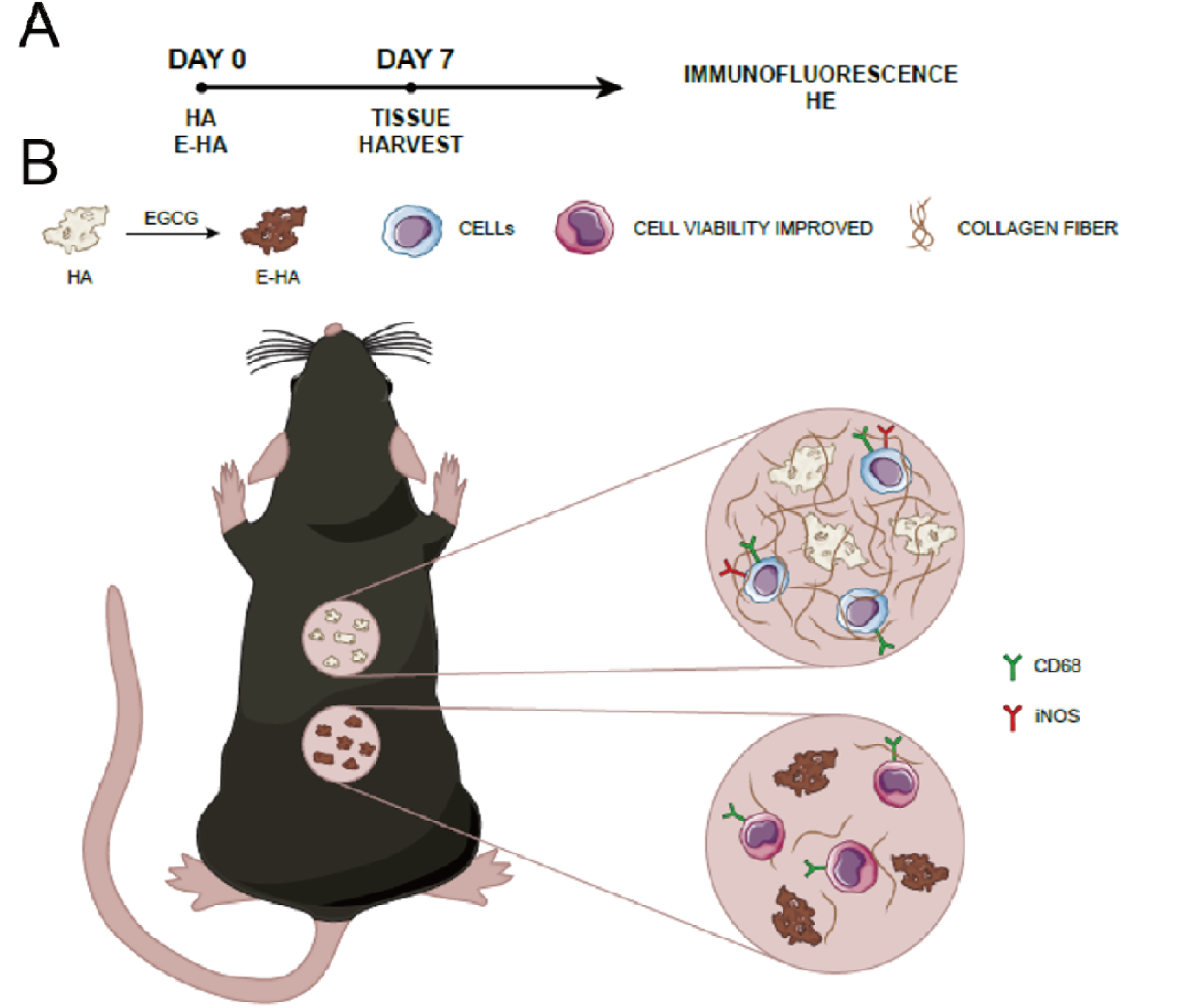
Schematic illustration. Diagram of *in vivo* experiments. (A) Showing the period to harvest and the subsequently executed testing. (B) Cell viability, macrophage polarization, and fibrosis outcome of EGCG modified bone graft, compared with bone graft alone.

## 2 Materials and Methods

### 2.1 Materials

We purchased commercial Heal-All^®^ Bone Repair Material from Zhenghai Biotechnology (Shandong, China), which contains primarily HA and collagen and maintains a three-dimensional porous structure. EGCG was obtained from Jiang Xi Lv Kang Natural Products (Jiang Xi, China). The solvents and chemicals used were all of analytical grade, with no further purification. To fabricate EGCG-modified HA for follow-up experiments, we dissolved EGCG in aseptic double-distilled water at a concentration of 0.64% (w/v). We then immersed HA in the EGCG mixture for 24 h at 4□ in the dark^14,20^. EGCG-modified bone graft (E-HA) was then washed with PBS three times and preserved in the dark at 4°C before use. The experimental process was conducted in accordance with the preservation protocol recommended by the manufacturer to avoid the oxidation denaturation of EGCG in light and heat. The optical density (OD) value of the EGCG solution at 272 was obtained before and after immersion to ensure successful fabrication of E-HA.

### 2.2 *In vitro* release profile of E-HA

E-HA was soaked in PBS in a 48-well plate at 37°C and 5% CO_2_ and scanned by ultraviolet–visible spectrophotometry to obtain the release profile. The OD value at 272 was used to detect EGCG in solution. At 0 h, the instrument was calibrated to zero; subsequently, we measured the OD value of the media at 2, 6, 12, 24, 48, 96, 120, 144, and 216 h. The media (200 μL) was removed and replenished with fresh buffer (200 μL) for each measurement. The solution concentration and EGCG weight were determined from an OD value for a specific concentration by establishing a standard curve with different concentrations (7.8, 125, 15.625, 31.25, 62.5, 125, 250, and 500 μg/mL); the vertical axis shows the absorbance value, and the horizontal axis shows the concentration of the EGCG solution. Linear regression analysis was performed using the weighted least-square method (W = 1/C2).

### 2.3 Surface morphology observation

A scanning electron microscope (SEM, s-800, HITACHI, Tokyo, Japan) operating at 15 kV and a digital camera (Canon EOS 6D Mark II) were used to characterize the morphology of the materials. A typical image of each sample was captured.

### 2.4 Cell viability

Murine RAW264.7 cells (RIKEN BioResource Center, Japan) were used for quantitative and qualitative cell viability assays. The cells were cultured in 48-well plates and seeded on untreated and EGCG-treated HA at 10^4^ cells/well cultured in 1640 medium supplemented with 10% fetal bovine serum. Immediately and after 1, 3, 5, and 7 days of culture, the cell viability was evaluated using Cell Counting Kit-8 (CCK-8, Dojindo Laboratories, Kumamoto, Japan) according to the manufacturer’s instructions. The OD value at 450 nm was measured by a micro-plate reader (Multiskan, Thermo, USA).

### 2.5 Establishment of an *in vivo* model

This experimental protocol was approved by the Institutional Review Board of West China Dental Hospital (No. WCHSIRB-D-2017-097). *In vivo* experiments were performed on 6-to 8-week-old C57 mice (Animal Experimental Center of Sichuan University) weighing approximately 37.5 g. Two parallel surgical sites were established on each mouse for pure HA and E-HA, respectively (N ≥ 3). To minimize the bias caused by the surgical site, the materials were implanted in different arrangements.

### 2.6 Surgical procedures

After being anesthetized with intraperitoneal injection 80 mg/kg ketamine and 16 mg/kg xylazine at a volume of 10 μL/g weight, the skin on the back of the mouse above the surgical area was shaved, and an aseptic process was applied before implantation. Two sagittal incisions of approximately 10 mm were made on the dorsal skin to create subcutaneous pockets for the bone repair material. Pure bone repair material and the EGCG-modified material were then implanted into the pockets. Subsequently, the incision was sutured with a horizontal mattress 3.0 Vicryl suture (Ethicon, CA, USA). The animals were housed in a room specially designed for animal experiments and were fed a standard laboratory diet. After 7 days of recovery, the animals were sacrificed. Then, bone repair material and whole skin samples around the surgical site were collected. All experiments were performed according to the guidelines of animal ethics committee of Sichuan university.

### 2.7 Hematoxylin-eosin (HE) staining

For HE staining, 8-μm sections were incubated at 65°C for 4 h to dewax. After ethanol dehydration, hematoxylin was applied for 5 min, and 1% hydrochloric acid ethanol was differentiated for 2 h. The sections were then incubated for 2 min in 0.2% ammonia water, followed by staining with eosin for 1 min. After dehydration, the sections were removed and fixed with neutral resin. The slides were viewed using a microscope with a 200 × oil immersion objective (Olympus Corporation, Tokyo, Japan) and scanned with a digital slide scanning system (PRECICE, Beijing, China).

### 2.8 Immunofluorescent staining

The paraffn-embedded tissue was cut into 8-μm-thick sections. After dewaxing and hydrating, immunostaining was conducted. Sections were pretreated with 0.1% Triton X-100 in PBS with 1% bovine serum albumin for 1 h, 1% t20 for 20 min, and then PBS for 20 min. Briefly, the sections were incubated for 30 min in the dark. The excess dye was washed away with PBS. Sections were incubated with isotype antibodies to exclude false positive staining. At least three parallel sections from different implantation sites were observed by fluorescence microscopy (Zeiss stereoscopic finding, V20, Germany). Five fields were randomly selected for immunofluorescence determination. Macrophages were magnified 400-fold to exclude false positive staining. Semi-quantitative analysis was then performed at a magnification of 40-fold. CaseViewer 2.1 and Image Pro Plus 7.0 (n = 5) were used to measure the fluorescence intensity of five random spots. Hoechst (ab138903, Abcam) staining analysis was performed at the same time as CD68 (ab222914, Abcam) and inducible nitric oxide synthase (iNOS, ab209027, Abcam) analysis.

### 2.9 Statistical analysis

All quantitative data are presented as the mean ± standard deviation. Statistical calculations were performed using GraphPad Prism 5.0 (San Diego, CA, USA). Semi-quantitative immunofluorescent staining data that did not conform to the normal distribution were further analyzed by the Mann–Whitney U test, while the statistical significance between groups was analyzed by analysis of variance, followed by Tukey’s multiple-comparison test. A value of P < 0.05 was considered statistically significant. A standard curve for the EGCG solution was established by linear regression. Pearson’s correlation analysis was used to analyze the detected concentration.

## 3 Results

### 3.1 *In vitro* release profile of E-HA

After HA was immersed in EGCG solution, the OD value decreased, indicating that the concentration had decreased and that E-HA had been successfully fabricated with EGCG adhering to the HA in the solution (**Fig. 3A**). The calculated concentration was based on the standard curve for the EGCG solution. To establish this standard curve, EGCG solutions were detected by ultraviolet–visible spectrophotometry, which revealed that EGCG has a maximum absorption wavelength of 272 nm, regardless of the concentration. Therefore, 272 nm was selected for detection, and the absorbance value was set as the vertical axis with the corresponding concentration set as the horizontal axis. A regression equation was obtained as y = 0.0067x–0.0327, with R^2^ = 0.9998. The linear concentration range was found to be 7.8–500 μg/mL for EGCG, as presented in **Fig. 3B**. A cumulative release profile of E-HA was obtained by detecting the amount of EGCG released to the buffer over time. With the obtained standard curve line, we can interpret the EGCG concentration corresponding to OD values measured at different time points in the cumulative release profile. The release profile demonstrated a minimal initial release, with only approximately 10 μg EGCG being released in the first 3 days, followed by a continuous release reaching approximately 100 μg over the next 3 days. An additional release of approximately 75 μg was measured at 216 h (**Fig. 3C**).

### 3.2 Surface morphology

The morphology of the bone repair material is shown in **Fig. 2A**. The morphology is inconsistent in shape, size, and porous features. After the addition of EGCG, the surface of the bone repair material became directional, dense and uniform. This change may be caused by hydrogen bonding between the EGCG and collagen fibers. The SEM results also show that the surface roughness decreased (**Fig. 2B**).

**Figure 2.**
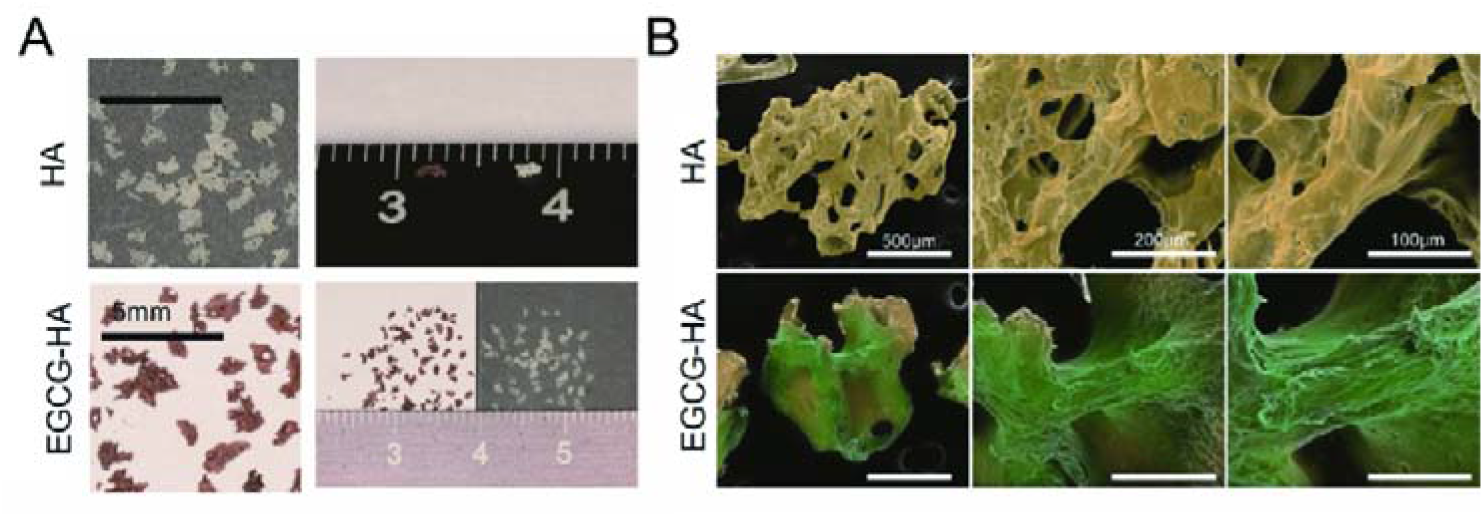
Representative image of HA and E-HA. (A) Surface morphology of materials is clearly presented under digital camera and (B) scanning electron microscope.

**Figure 3.**
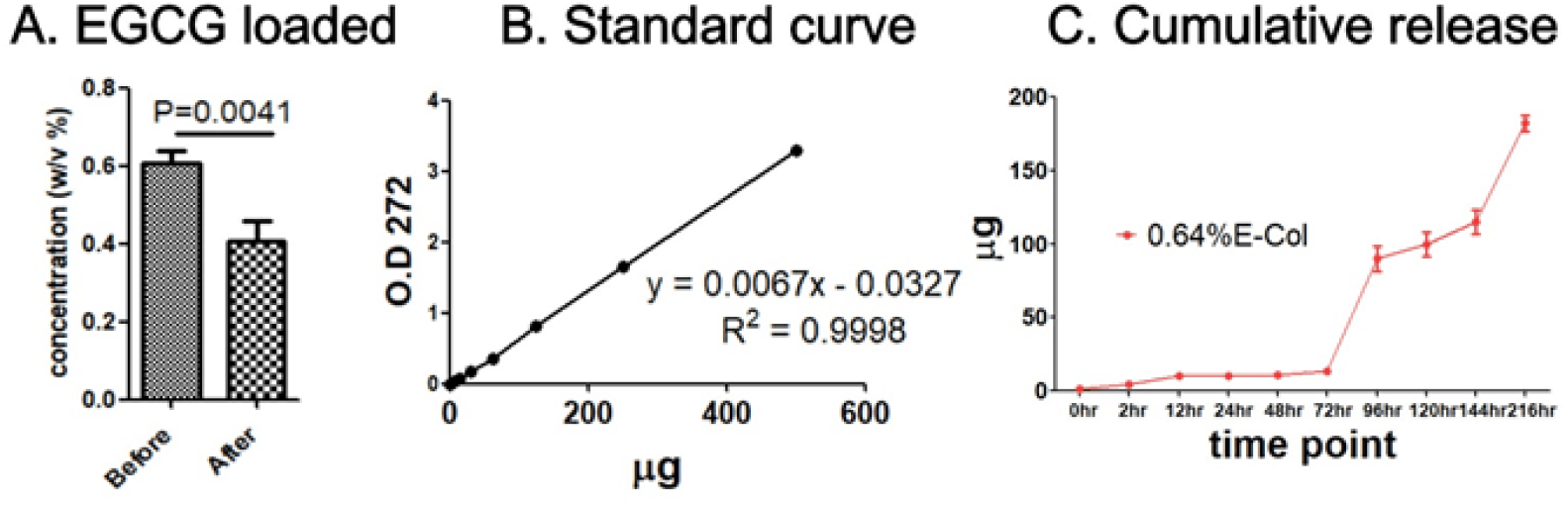
The establishment and release of EGCG on the bone graft. **(A)** E-HA fabrication assessment by detecting the concentration of EGCG solution before and after immersing HA. (B) The standard curve of EGCG solution at the concentration between 7.8 and 500 μg/ml, (C) obtained under OD value of 272. *In vitro* cumulative release profile of E-HA.

### 3.3 Cell viability

RAW 264.7 cells were seeded on the material for 7 days; throughout this period, the cell viability remained relatively constant for the HA group, whereas the effect of EGCG on cell viability increased as the co-culture time increased. On days 0 and 1, there was no significant difference between the HA and E-HA groups; however, on days 3 and 5, the cell viability improved slightly in the group with EGCG. The viability continued to increase, reaching a maximum on day 7 (**Fig. 4**). Thus, the modification of EGCG did not negatively impact cell viability in RAW 264.7 cells; rather, EGCG appears to improve cell viability.

**Figure 4.**
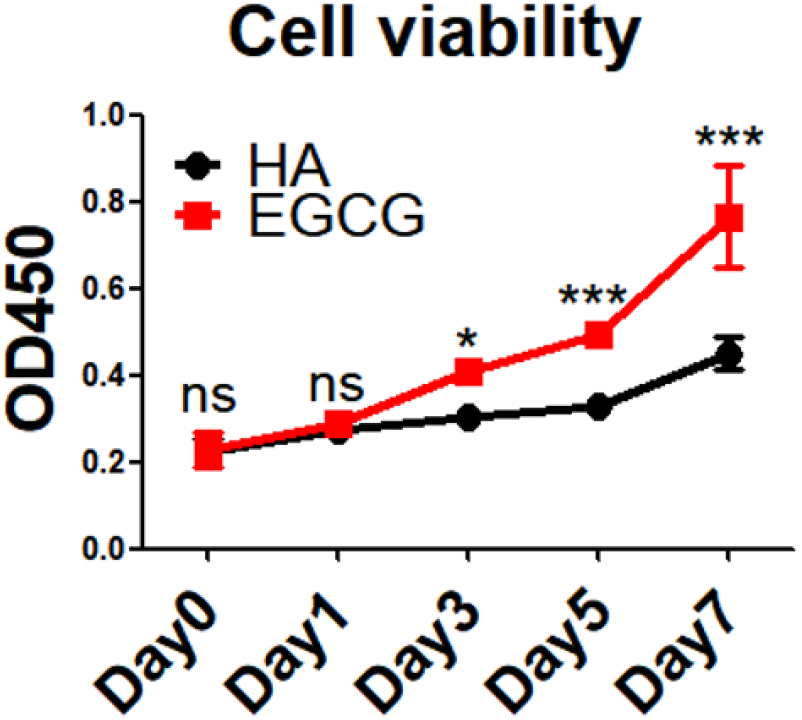
Cell viability assay. Culturing RAW 264.7 on deproteinized bone material with or without EGCG treatment, then the cell viability was detected on day 0, 1, 3, 5 and 7. NS= no significance; *= P<0.05, **= P<0.01, ***= p<0.001.

### 3.4 Fibrosis severity after HA and E-HA implantation in relation to macrophage phenotype

To determine whether E-HA can mitigate fibrosis by modulating the immune microenvironment, the selected materials were implanted subcutaneously (**Fig. 5A–F**), and the reaction of immune cells towards EGCG modification was investigated by HE staining (**Fig. 5G**). Within 7 days after implantation of the bone repair material, the incisions of both groups had healed well. A nonvascular tubular structure was observed, which was more abundant in the E-HA group. The fibrous encapsulation effect was much more intense in the HA group, with a large amount of collagen fibers deposited on the surface of the material. The surrounding immune cells migrated to the bone repair material, forming an inflammatory cell infiltration zone. This inflammation resulted from the implantation of EGCG-loaded bone repair material, which is lighter than pure bone repair material. We further explored the macrophage phenotype at this time point. Based on Hoechst, CD68, and iNOS immunofluorescence staining images (**Fig. 5H**), we found that the two groups recruited macrophages of different phenotypes. CD68, which serves as a surface marker for macrophages, was observed in both groups. However, iNOS, an indicator of pro-inflammatory phenotype macrophages, was not detected in the E-HA group, although it was detected in the HA group, reconfirming macrophage infiltration and the effect of EGCG on anti-inflammation.

**Figure 5.**
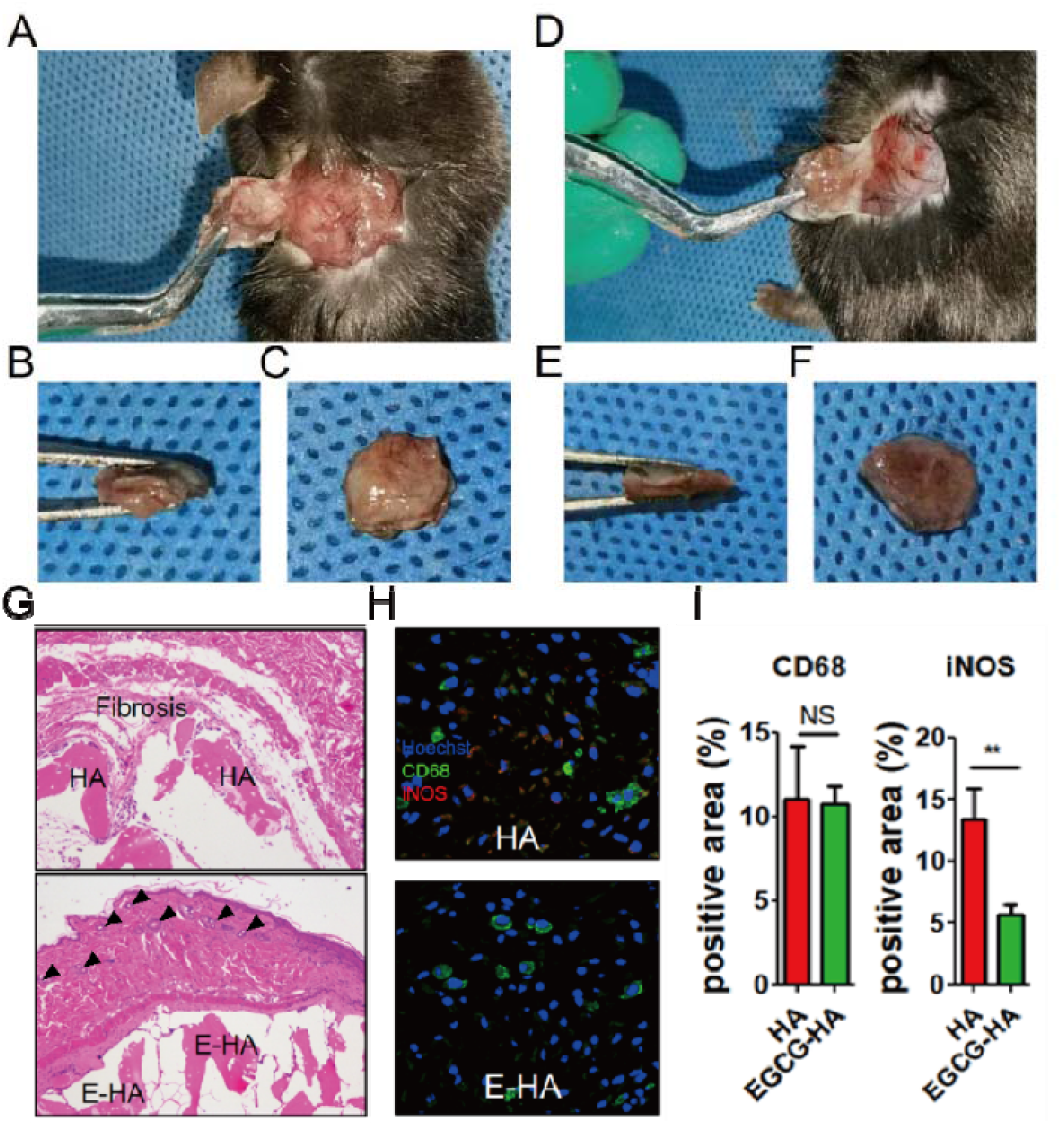
Evaluation of fibrosis and phenotype of infiltrating macrophages after implantation. (A-C) Pictures of harvested tissue encapsulated with (D-E) HA and E-HA. (G) HE staining result on day 7 post-surgery. (H) Immunofluorescent staining result on day 7 postsurgery. Hoechst, blue; CD68, green; iNOS, red. (I) Semi-quantitative analysis of immunofluorescent staining result. Arrows indicate nonvascular tubular structure. NS= no significance; **= P<0.01.

## 4 Discussion

Serious defects in hard tissue can inhibit self-healing, with critical-sized bone defects requiring surgical intervention^21^. At the present, biomaterials and GBR membranes are often used in clinical practice to block the invasion of soft tissue in the growth space of the hard tissue, a practice that has been widely validated. Among these materials, collagen membranes modified by EGCG have been thoroughly investigated, including studies on their biotoxicity^22^, effect on osteoblasts^20^, chemotaxis of macrophages, phenotypic regulation, and FBR. The ideal therapeutic effect of an implanted biomaterial cannot be achieved if the stimulated effect due to modification is neglected. For example, biomaterials often include HA, such as BMP-2-immobilized PLGA/hydroxyapatite fibrous scaffold^25^, or have been modified, for example, by loading nerve growth factor^26^. Therefore, to better stimulate the bone ECM and to provide a suitable microenvironment for the regeneration of hard tissue, a bovine-derived decellularized bone graft composed of HA and collagen type I, which is similar to the composition of bone and current commercial dental bone substitutes, was adopted. When a biomaterial is implanted, osteoblasts are involved in the repair, and the interaction between the material and immune cells is crucial in the tissue healing process. Protein adsorption is the beginning of the acute inflammatory phase of FBR, and the inflammation will gradually transit toward tissue repair. Immediately after surgery, macrophages quickly identify the implant, are recruited to the biomaterial, and form a specific immune microenvironment. The participation of macrophages and T cells in the reconstruction of ECM after *in vivo* implantation is highly important for osteogenesis^27^. Macrophages are involved in the initiation of inflammation and the following tissue reconstruction due to the secretion of cytokines. A transformation in the macrophage phenotype from pro-inflammatory to anti-inflammatory is necessary in both hard- and soft-tissue repair. Long-term or severe inflammation usually contributes to fibrous capsule formation by activating anti-inflammatory macrophages releasing fibrotic cytokines, which block the surrounding material and eventually lead to bone-biomaterial integration failure^28^. As one of the primary causes of implant failure, aseptic loosening is caused by the formation of wear particles from the implant under phagocytosis, which promotes macrophage secretion of cytokines related to bone absorption^29^. HA-induced phagocytosis depends on IL-1β and TNF-α produced by macrophages^29^, which are closely related to the pro-inflammatory phenotype. Furthermore, a long-term high pro-inflammatory/anti-inflammatory phenotypic ratio of macrophages often indicates implantation failure. Thus, in addition to manufacturing biomaterials similar to the natural ECM, the phenotype of macrophages should also be considered. A biodegradable three-dimensional scaffold comprising polymer matrix (polylactic acid and polyethylene glycol), nano-HA, and dexamethasone has been shown to promote macrophage polarization towards the anti-inflammatory phenotype and osteogenesis, in which pro-inflammatory macrophages downregulate IL-6 and iNOS expression. In fact, both pro-inflammatory and anti-inflammatory phenotypes are indispensable for tissue healing, and pro-inflammatory macrophages can even guide the subsequent behavior of anti-inflammatory macrophages. However, their temporal behavior is important, as successful repair can never be accompanied by excessive pro-inflammatory polarization. As CD68 is widely acknowledged as a marker for macrophages, CD staining was applied to identify the presence of macrophages, and we confirmed the reduction of pro-inflammatory phenotypes due to EGCG addition based on iNOS. Apart from inhibiting pro-inflammatory macrophages, we also found that EGCG alters the surface morphology, as shown by SEM. The morphology is closely related to FBR. Researchers have speculated that the mechanism by which HA coatings promote osteogenesis may be related to the surface morphology rather than the stimulation of macrophages to express BMP-2. HA can significantly up-regulate pro-inflammatory factors, such as TNF-α, in pro-inflammatory macrophages and can further inhibit anti-inflammatory macrophages from expressing BMP-2, as occurs in osteoconductive condition^32^. Thus, HA has a pro-inflammatory tendency and is not conductive to the expression of BMP-2; moreover, both of these effects may be bolstered by the addition of EGCG^9,12^. However, further investigation is needed to determine whether EGCG can promote the polarization of macrophages toward anti-inflammatory phenotypes and stimulate the secretion of desired cytokines, as demonstrated by our previous studies on collagen membranes. Cytotoxicity is another essential aspect to evaluate when considering biocompatibility. Karin H. Müller and Michael Motskin *et al*. reported that HA nanoparticles form large agglomerates in media, resulting in extensive particle uptake in macrophages and sequestration in surface-connected compartment^33^, triggering a dosedependent cytotoxicity. To reduce the cytotoxicity, the particles can be citrated or dispersant Darvan 7 can be added^34^. Our HA particle size was 1000–2000 μm, and the cell viability was improved by the addition of EGCG. We found that the effect of EGCG on cell viability was related to its release from E-HA. Our *in vitro* EGCG release rate assay showed that more EGCG was released over time. As the culture time increased, the effect of EGCG on cell viability grew. Therefore, it is suggested that the addition of EGCG will not reduce, and may even increase, cell activity, which is beneficial for healing. In this article, we considered only a subcutaneous method for reducing pro-inflammatory macrophages. The specific mechanism and phenotypic transformation, including the ratio of pro-inflammatory/anti-inflammatory macrophages at different time points in a bone model will be discussed in the future.

## 5 Conclusion

In summary, we have presented a potential strategy for modifying bone repair material. This strategy provides an anti-inflammatory effect by reducing the accumulation of pro-inflammatory macrophages and further regulates fibrosis, although the effect of EGCG on the surface morphology cannot be excluded. The specific mechanism leading to reduced fibrosis requires verification. The results of this study will pave the way for subsequent experiments and for further work on the osteogenesis effect of E-HA, as this is a essential effect of bone repair materials. Under the studied circumstances, the addition of EGCG to HA improved cell activity, and no CD68^+^iNOS^+^ cells were detected by immunofluorescence staining at 7 days post-surgery, indicating that the addition of EGCG may resolve the following regenerative cascade.

## Data Availability Statement

No datasets were generated or analyzed during the current study

